# Ribovirus classification by a polymerase barcode sequence

**DOI:** 10.1101/2021.03.02.433648

**Authors:** Artem Babaian, Robert C. Edgar

## Abstract

RNA viruses encoding a polymerase gene (riboviruses) dominate the known eukaryotic virome. Next-generation sequencing is revealing a wealth of new riboviruses with uncharacterised phenotypes, precluding classification by traditional taxonomic methods. These are often classified on the basis of polymerase sequence identity, but standardised methods to support this approach are currently lacking. To address this need, we describe the polymerase palmprint, a well-defined segment of the palm sub-domain delineated by well-conserved catalytic motifs. We present a novel algorithm, `Palmscan`, which identifies palmprints in nucleotide and amino acid sequences. We describe PALMdb, a reference database of palmprints derived from public sequence databases. `Palmscan` source code and PALMdb data are deposited at https://github.com/rcedgar/palmscan and https://github.com/rcedgar/palmdb, respectively.

## Introduction

### Background

Metagenomic sequencing is driving an exponential growth in the number of known microbes, where the deluge of new data and lack of observed phenotype other than sequence preclude traditional taxonomic methods based on expert review of traits such as virion size and shape. This has prompted the development of new approaches based on so-called marker, tag, or barcode regions of microbial genomes. Microbial barcode databases, including SILVA (Pruesse et al. 2007) for ribosomal small-subunit RNA and UNITE (Abarenkov et al. 2010) for the fungal internal transcribed spacer region, have been developed to support this approach.

There are millions of extant viral species (Geoghegan and Holmes 2017), but less than five thousand are currently recognised by the International Committee on Taxonomy of Viruses (ICTV 2019a). In virology, classification on the basis of marker genes, typically polymerases, is well established (Koonin et al. 2020), but progress is impeded by a lack of standardised sequence analysis methods and databases comparable to SILVA and UNITE.

In this work, we developed methods for barcode-based classification of *Riboviria*, a group which includes all RNA viruses encoding an polymerase gene, either RNA-dependent RNA polymerase (RdRP) or RNA-dependent DNA polymerase (reverse transcriptase, RT). Here, we use the term polymerase to refer to viral RdRP or viral RT unless otherwise stated. Riboviruses account for 35% of named virus species in the NCBI Taxonomy database (Federhen 2012), infecting a wide variety of animals and plants (Garcia-Villada and Drake 2012). We show that a ∼100aa sub-sequence of the polymerase gene, which we call the palmprint, is well-suited to automated analysis. The palmprint is delineated by well-conserved motifs in the polymerase palm sub-domain which are essential to polymerase function. We describe a novel algorithm, `Palmscan`, to identify palmprints in longer sequences such as virus genomes and ORFs. We present a reference database of palmprints, PALMdb.

### Polymerase gene

The polymerase fold belongs to the template-dependent nucleic acid polymerase superfamily; it resembles a grasping right hand with thumb contacting finger (Mönttinen et al. 2014). Surface regions directly involved in nucleotide selection or catalysis are strongly conserved, in particular short motifs conventionally designated by letters A through G found in the active site. Motifs A, B and C are found in the palm sub-domain and are well conserved in most known RdRPs (Velthuis 2014). A and C contain essential aspartic acid residues which coordinate the Mg+/Mn+ cation for catalysing phosphodiester bond formation, while B contains an almost perfectly conserved glycine required for nucleotide selection. The motifs appear in ABC (canonical) order in the primary sequence of most known polymerases, but the active site sequence is permuted into CAB order in several independent lineages (Gorbalenya et al. 2002; Sabanadzovic, Abou Ghanem-Sabanadzovic, and Gorbalenya 2009); see Fig. 1. Amino acid identity can be as low as 10% between diverged species (Bruenn 2003). The boundary of the polymerase domain may be unclear as the gene may be embedded in a longer open reading frame (ORF) together with other domains. Significant sequence similarity with a known viral polymerase is not sufficient to establish viral origin or polymerase function. Viral RdRPs can be integrated into host germline DNA via reverse transcription, becoming an endogenous viral element (EVE) (Holmes 2011; Feschotte and Gilbert 2012). Viral RTs are ubiquitously inserted into host germlines, losing function over time and leaving “fossils” with recognisable sequence similarity (Bock and Stoye 2000).

**Figure 1.**
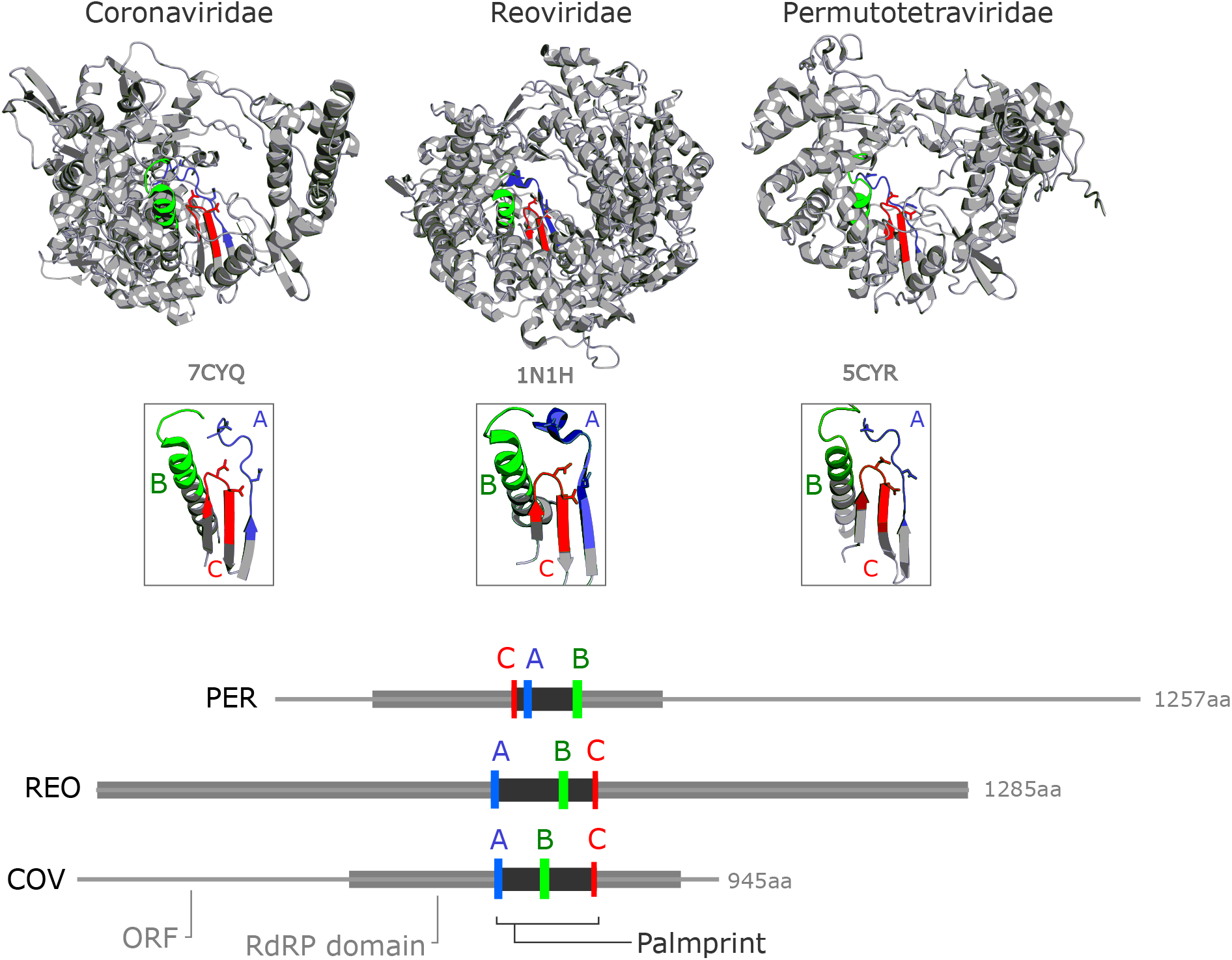
RdRP gene structure. Structural conservation of catalytic motifs among three divergent RdRP structures in the Protein Data Bank (Berman et al. 2000). *Coronaviridae* (COV, virus: SARS-CoV-2, pdb:`7CYQ`.) is from a positive-strand RNA virus. *Reoviridae* (REO, virus: Mammalian orthoreovirus 3 Dearing, pdb:`1N1H`) is from a double-stranded RNA virus, and *Permutotetraviridae* (PER, virus: Thosea asigna virus, pdb:`5CYR`) is a positive-stranded RNA virus with permutation of motif C. RdRP domains defined by PFAM (COV: `RdRP 1`, TAV: `DNA/RNA pol sf`, and REO: `RdRP 5`) are shown within the open reading frame for REO and PER, and the mature polyprotein cleavage peptide from the 7,096 aa ORF1ab for COV.

### Palmprint

We defined the polymerase palmprint to be the minimal segment containing all letters of the A, B and C motifs (Figs. 1 and 2). In domains where the motifs appear in canonical order, the palmprint extends from the first letter of A to the last letter of C; in CAB permuted domains it extends from the first letter of C to the last letter of B. To extract palmprints from longer sequences, we implemented an algorithm, `Palmscan`, to identify A, B and C motifs using position-specific scoring matrices, (Stormo et al. 1982)), described in Methods.

**Figure 2.**
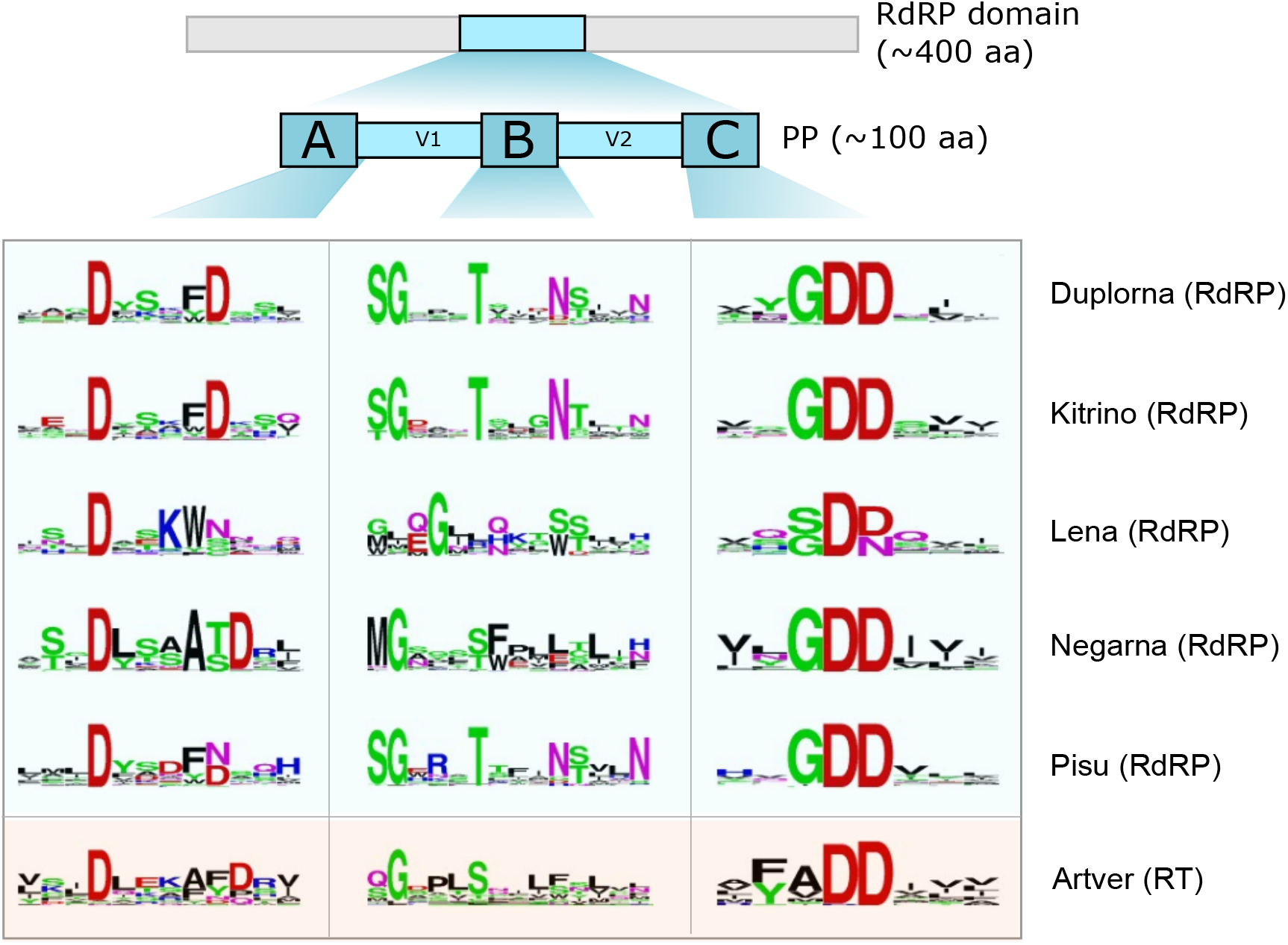
Polymerase palmprint segment with motif logos. The palmprint segment is a ∼100aa region in the active site of the polymerase domain. Motifs A, B and C are well-conserved. The figure shows sequence logos for the five RdRP-containing ribovirus phyla (Duplorna=*Duplornaviricota*, etc.) and one for RTs in *Artverviricota*.

**Figure 3.**
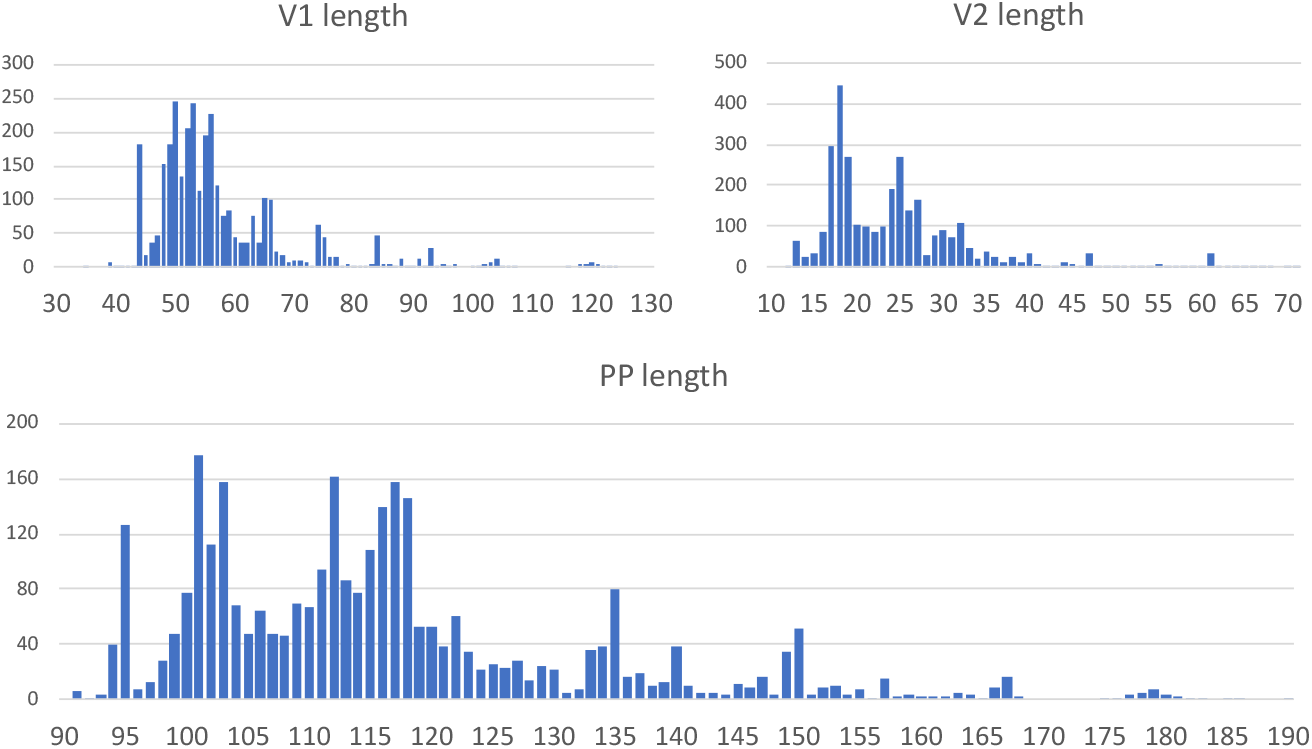
Lengths of the V1 and V2 variable regions and palmprint segment. Distributions were measured on full-length RefSeq Ribovirus genomes.

**Figure 4.**
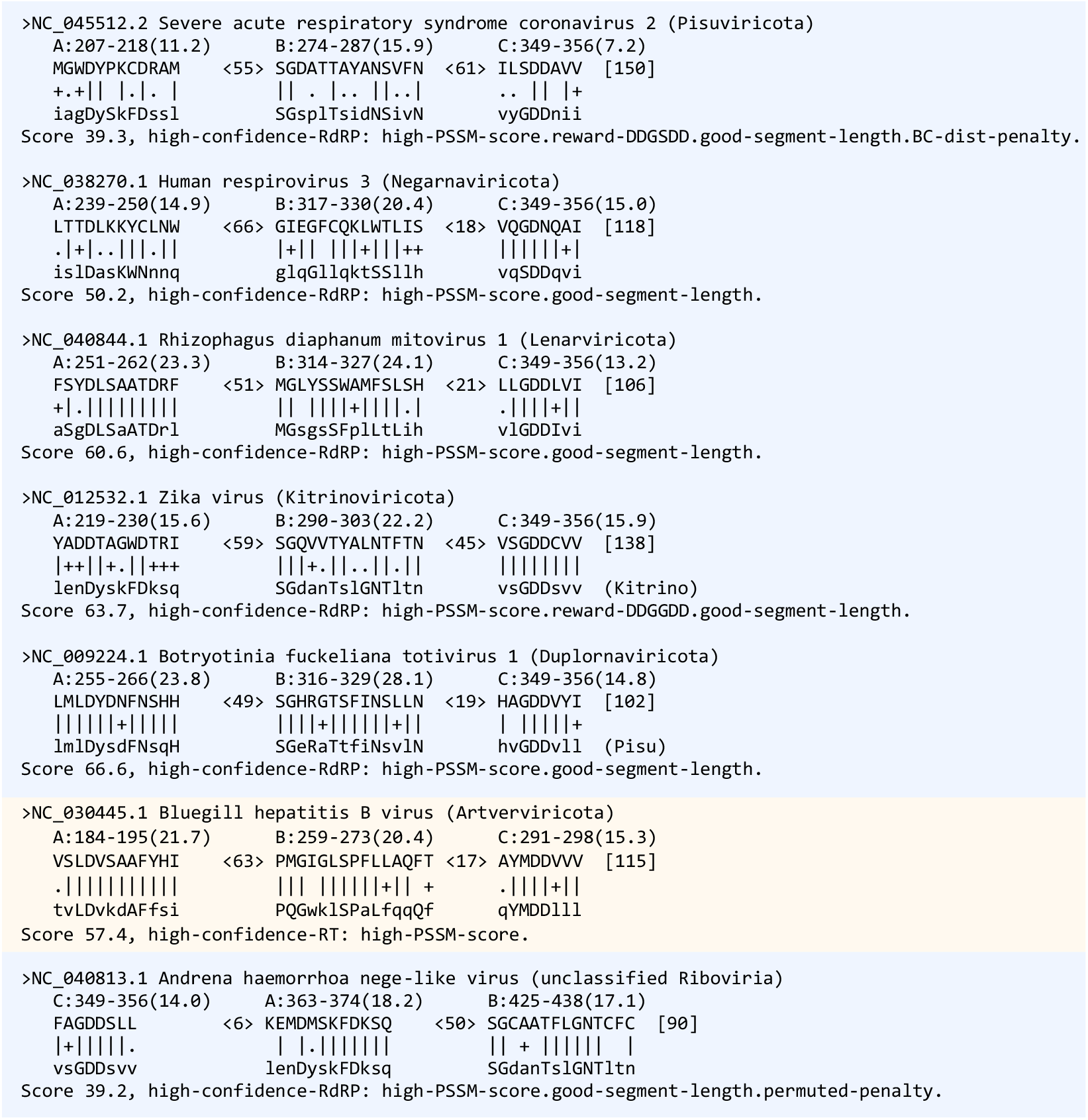
Example Palmscan report. Report showing motif alignments for RefSeqs in each phylum, plus one unclassified genome (`NC 040813.1`) with permuted motifs. See Fig. 5 for explanation of information shown in the report.

**Figure 5.**
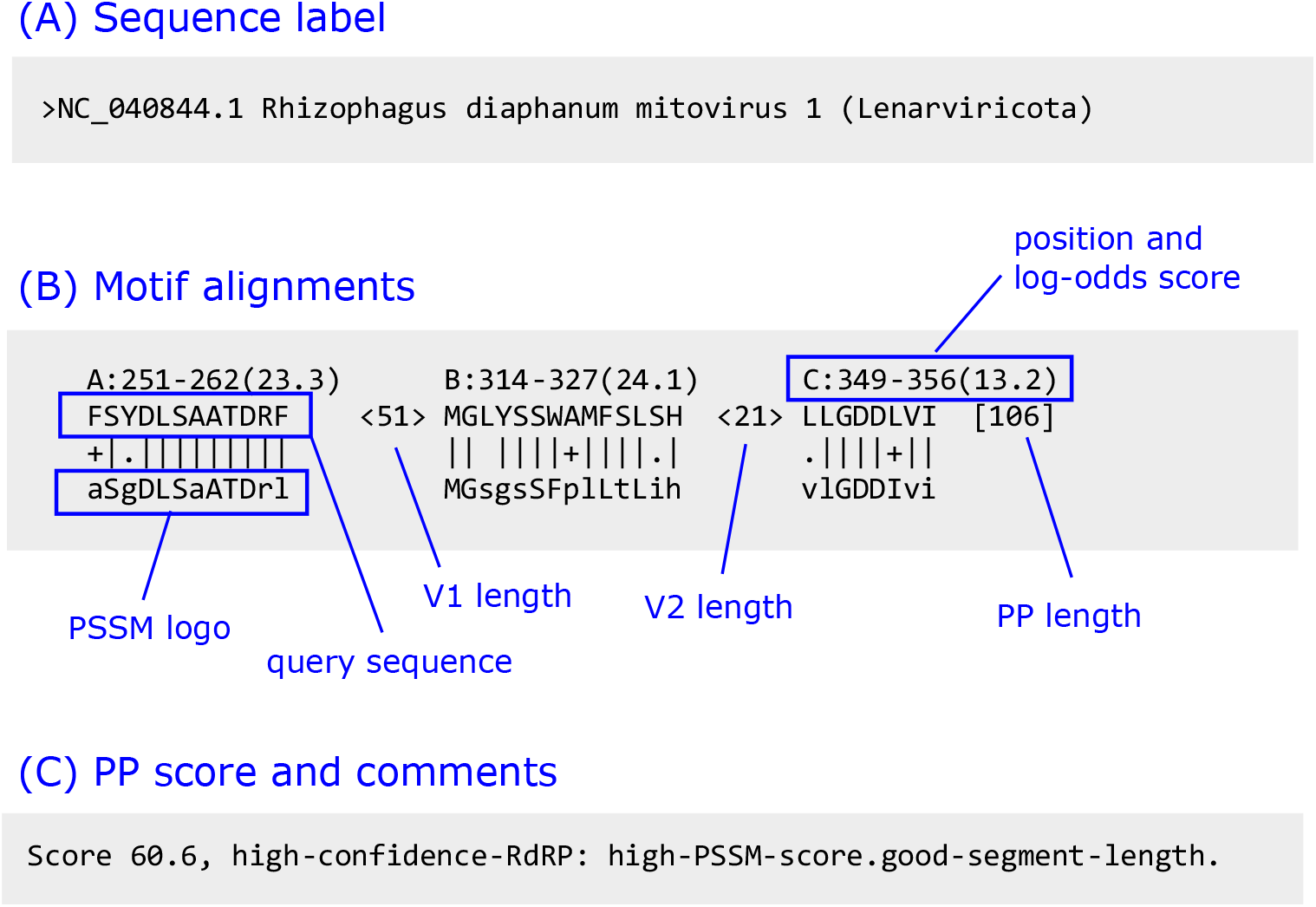
Annotated Palmscan alignment. An alignment report comprises (A) query label from FASTA input file, (B) motif alignments and lengths of V1, V2 and palmprint regions, and (C) final score and comments.

### Sequence-based classification

Several approaches have been proposed for classifying microbial barcodes. SILVA constructs a predicted phylogenetic tree for 16S ribosomal RNA sequences. Placement of metagenomic sequences in the tree is compared with consensus subtrees for named groups to infer membership of taxonomic clades, often leaving lower ranks such as species and genus unclassified, reflecting the difficulty of automated taxonomy prediction and the large number of groups known only from sequence which are not yet named. Alignment and tree construction of large numbers of highly diverse sequences is challenging, and inferences from the resultant trees are controversial (Edgar 2018a; Holmes and Duchêne 2019). RDP (Maidak et al. 1996) uses a Naive Bayesian Classifier (NBC) based on nucleotide 8-mer frequencies to predict taxonomy (Wang et al. 2007). The NBC does not require a tree, but also leaves lower ranks unclassified when sequences do not belong to named species. Both tree-based and NBC methods may give different classifications to a given sequence as the database expands. UNITE constructs operational taxonomic units (OTUs) (Sneath, Sokal, et al. 1973) by clustering ITS sequences at 99% identity, identifying clusters with named clades where possible and otherwise designating OTUs as so-called species hypotheses (SHs). SH identifiers are designed to provide a stable reference system so that newly-named groups can be associated with SHs, and metagenomic sequences will be associated with the same SH identifier as the database expands. Our approach in this work is similar to UNITE’s: we circumvent challenges with multiple alignment, tree inference and database growth by clustering palmprints into stable species-like OTUs.

### OTU construction

Many methods for constructing OTUs from barcode sequences have been proposed, notably agglomerative clustering (Schloss and Handelsman 2005) and centroid clustering (Edgar 2013), with competing and contradictory claims that some methods are better than others (see (Edgar 2018b) and references therein). Our view, following the conclusions of (Edgar 2018b), is that most clustering methods can generate equally good OTUs providing that high-quality sequences are provided as input and thresholds are selected appropriately. For this work, we designed an OTU pipeline with the following goals: computational efficiency, good agreement with taxonomy, and stability.

## Methods

### Position Specific Scoring Matrices

We created position-specific scoring matrices (PSSMs) (Stormo et al. 1982) to recognise A, B and C motifs. Intuitively, a PSSM is a numerical representation of a sequence logo. More formally, a PSSM *P* [*i, j*], *i* = 1 … *L, j* = 1 … 20 represents a multiple alignment of length *L* columns by a log-odds score for each possible amino acid in every column. Letters found with high frequency at a given position are assigned positive scores, while unobserved and low-frequency letters have negative scores. The score of *P*_*i*_ aligned to the *k*th letter in a query sequence *Q* is *P* [*i, Q*_*k*_], and the total score of the alignment is the sum of scores

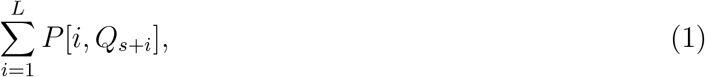

where *s* is the starting position in *Q*. Gaps are not permitted in the alignment. Log-odds scores were calculated from multiple sequence alignments (MSAs) for each motif assuming background amino acid frequencies from BLOSUM62 (S. Henikoff and J. G. Henikoff 1992) with pseudo-count frequency 0.1. Seven sets of PSSMs were constructed: one for each phylum where RdRP is commonly found, i.e. *Duplornaviricota, Kitrinoviricota, Lenaviricota, Negarnaviricota* and *Pisuviricota*; one for reverse transcriptases in *Artverviricota*; and one for permuted members of the *Birnaviridae* family, which have distinctly different motifs. MSAs for each motif in each set were constructed as follows. First, a small seed alignment was constructed by hand. The resulting PSSM was then used to search polymerase-containing datasets. This process was iterated until no further improvement was obtained. This may have resulted in some degree of over-training to known polymerases, but sensitivity was not our primary goal; we preferred to ensure that positive PSSM alignment scores were strongly predictive of a valid viral motif.

### Segment length distributions

In a palmprint with canonical ABC motif order, we define the V1 segment to be the sequence between A and B; similarly V2 is the sequence between B and C. In a permuted palmprint with CAB order, V1 is the sequence between C and A; V2 is the sequence between A and B. The V1 and V2 (variable) segments are less well conserved than the motifs. The lengths of the V1 and V2 segments and the total length of the palmprint vary within constraints imposed by structure and function; length distributions were measured for ribovirus RefSeqs.

### Palmscan algorithm

We implemented an algorithm, `Palmscan`, to identify palmprints in longer sequences and discriminate RdRPs from RTs. The first step is to align all PSSMs to all positions in the query sequence, which is six-frame translated in the case of a nucleotide query. The highest-scoring alignment for each PSSM is recorded, and the motif set with highest total log-odds score is identified. If the order is not ABC or CAB, the query is rejected. Otherwise, the lengths of V1, V2 and putative palmprint are determined. A palmprint score is derived, starting with the sum of log-odds scores of the three PSSMs in the highest-scoring set, adjusted by a series of heuristics which are designed to quantify qualitative judgements that might be made by a human expert.

For example, if any PSSM log-odds score is < 2 then the alignment is rejected; if the length of V1 is less than 35aa then a penalty of 5 is subtracted from the score. These heuristics were developed through an iterative process of manual review and improvement of `Palmscan` predictions on ribovirus RefSeqs and decoys. If the final score is ≥ 20, the palmprint is reported.

### Construction of reference palmprints

We used the following datasets in validation and training. `Wolf18` is the `RNAvirome.S2.afa` file from (Wolf, Kazlauskas, et al. 2018). `PF00680 RdRP 1`, `PF00978 RdRP 2`, `PF00998 RdRP 3` and `PF02123 RdRP 4` are full alignments of PFAM (Bateman et al. 2004) RdRP models; similarly `PF00078 RT1` and `PF07727 RT2` are PFAM RT alignments. `Genomes` is the set of RefSeq complete genomes from ribovirus phyla that typically contain RdRP (see Introduction). `GB241nt` is all nucleotide sequences from GenBank (Benson et al. 2012) v241, `GB241aa` is all translated CDSs from GenBank v241, `NR` is the NCBI non-redundant protein database (Pruitt, Tatusova, and Maglott 2005). `Palmscan` hits from all these sources were combined into a set of reference sequences, excluding those matching RTs. Those annotated by the source database as non-viral were retained to support discrimination of viral RdRP palmprints from RTs and non-viral palmprint-like sequences.

### Decoy set of non-RdRP sequences

We created a consolidated set of non-RdRP sequences (`Decoy`) to aid the development and validation of `Palmscan`, including the following amino acid sequences clustered at 97% identity: UniProt proteomes for human (`UP000005640`), yeast (`UP000002311`), *E. coli* (`UP000000558`); all retroviral and DNA-viral sequences from GenBank; `PF00078 RT1` and `PF07727 RT2`; the training set of validated RT developed for `myRT` (Sharifi and Ye 2021), and disordered proteins from https://disprot.org (Hatos et al. 2020). Manual inspection of the following ten sequences identified as high-confidence RdRP by `Palmscan` were confirmed by InterPro (Hunter et al. 2009) and `BLAST` (Altschul et al. 1997) to be RdRP and discarded: `CZQ50745.1`, `AFH02745.1`, `AFH02746.1`, `ADO67072.1`, `pdb|4TN2|A`, `YP 009506261.1`, `ADE61677.1`, `YP 009551602.1`, `AXY66749.1`, and `AWS06671.1`.

### OTU clustering

We chose to construct OTUs using the `UCLUST` algorithm (Edgar 2010) as implemented in the `cluster fast` command of `USEARCH`. The objective of `UCLUST` at a given identity threshold *t* is to identify a set of centroid sequences such that (1) all centroids have identity < *t* to each other, and (2) all other sequences in the input have identity ≥ *t* to a centroid (Fig. 6). Typically, many different solutions to this objective exist for a given set of input sequences, and this can be exploited to enable a stable update procedure, i.e. a method for adding new sequences without changing existing cluster assignments that satisfies the objective after updating. Given an existing set of centroids *C*, new sequences are aligned to *C* and those having identities ≥ *t* are assigned to existing clusters. The remainder are clustered by `UCLUST`, giving new centroids which are added to *C*.

**Figure 6.**
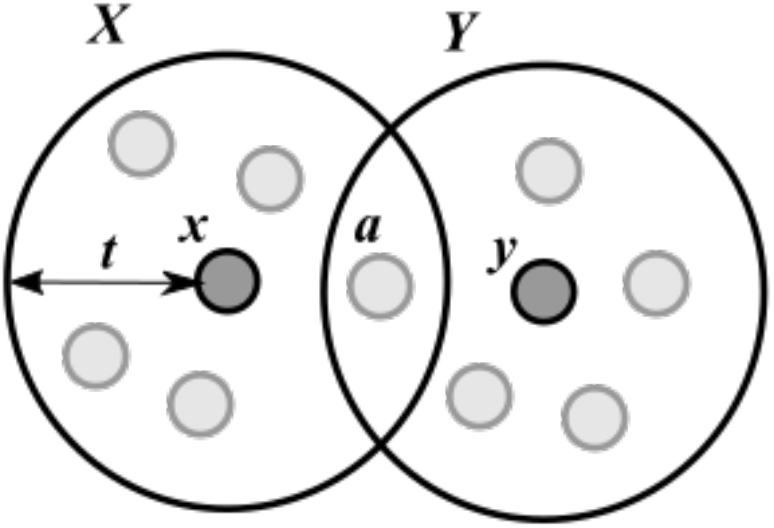
Centroid clustering. The objective of centroid clustering is to identify a set of centroid sequences such that all centroids are less than identity *t* to each other, and all other sequences are ≥ *t* to a centroid. The figure shows two clusters *X* and *Y* defined by centroids *x* and *y* respectively. Dots represent sequences; distances indicate identity. The identity threshold *t* can be considered the radius of a cluster. Some sequences may have identity ≥ *t* to two or more clusters, in which case they are assigned to the centroid with highest identity. There may be ties for highest identity, as for *a* in the figure, in which case the assignment is chosen arbitrarily.

### Taxonomy annotation of reference palmprints

We constructed a set of named reference sequences by selecting all palmprints having at least one taxon name in ranks we consider to be canonical: phylum, class, order, family, genus and species, keeping only names at those ranks. For example, `NCBI:txid1532164` is `superkingdom:viruses`, `clade:Riboviria`, `kingdom:Orthornavirae`, `phylum:Pisuviricota`, `class:Pisoniviricetes`, `order:Nidovirales`, `suborder:Cornidovirineae`, `family:Coronaviridae`, `subfamily:Orthocoronavirinae`, `genus:Alphacoronavirus`, `no rank:unclassified Alphacoronavirus`. In our named reference set, a sequence with `NCBI:txid1532164` is annotated as `phylum:Pisuviricota`, `class:Pisoniviricetes`, `order:Nidovirales`, `family:Coronaviridae`, `genus:Alphacoronavirus`. In this example, non-canonical ranks including suborder and subfamily are omitted. Species is not named and is therefore omitted; other canonical ranks are named and included.

### Local alignment methods

We used `diamond blastx --masking 0 --mid-sensitive -s 1 -c1 -p1 -k1 -b 0.75` (v2.0.7) (Buchfink, Xie, and Huson 2015) and `BLASTP` with default settings to create local alignments and E-values.

### Prediction accuracy metrics

We use positive predictive value (PPV) and false discovery rate (FDR) as our accuracy metrics for palmprint identification and classification algorithms. PPV is the number of correct predictions divided by the total number of predictions; FDR is the number of incorrect predictions divided by the total number of predictions. By design, queries which are unclassified by a prediction method are not considered by these metrics.

### Expect value of Palmscan false positives

The expect value, or E-value, of a search algorithm is the expected number of false positive detections at or above a given score in a random sequence of the same size as the search space. For `Palmscan`, we measured the number *F* of reported palmprints with score ≥ 20 in random sequences of 10^11^ nucleotides and amino acids, respectively. The E-value for a search space of length *L* letters, i.e. the number of false-positive palmprints expected by chance, is then *FL/*10^11^. None of the false-positive sequences had `blastp` E-values < 1 against a database of RefSeq ribovirus polymerases. We therefore recommend using `blastp` to confirm `Palmscan` predictions.

## Results

### Score threshold tuning

We set the minimum score to report a high-confidence RdRP palmprint to 20 after review of the score distributions shown in Fig. 7.

**Figure 7.**
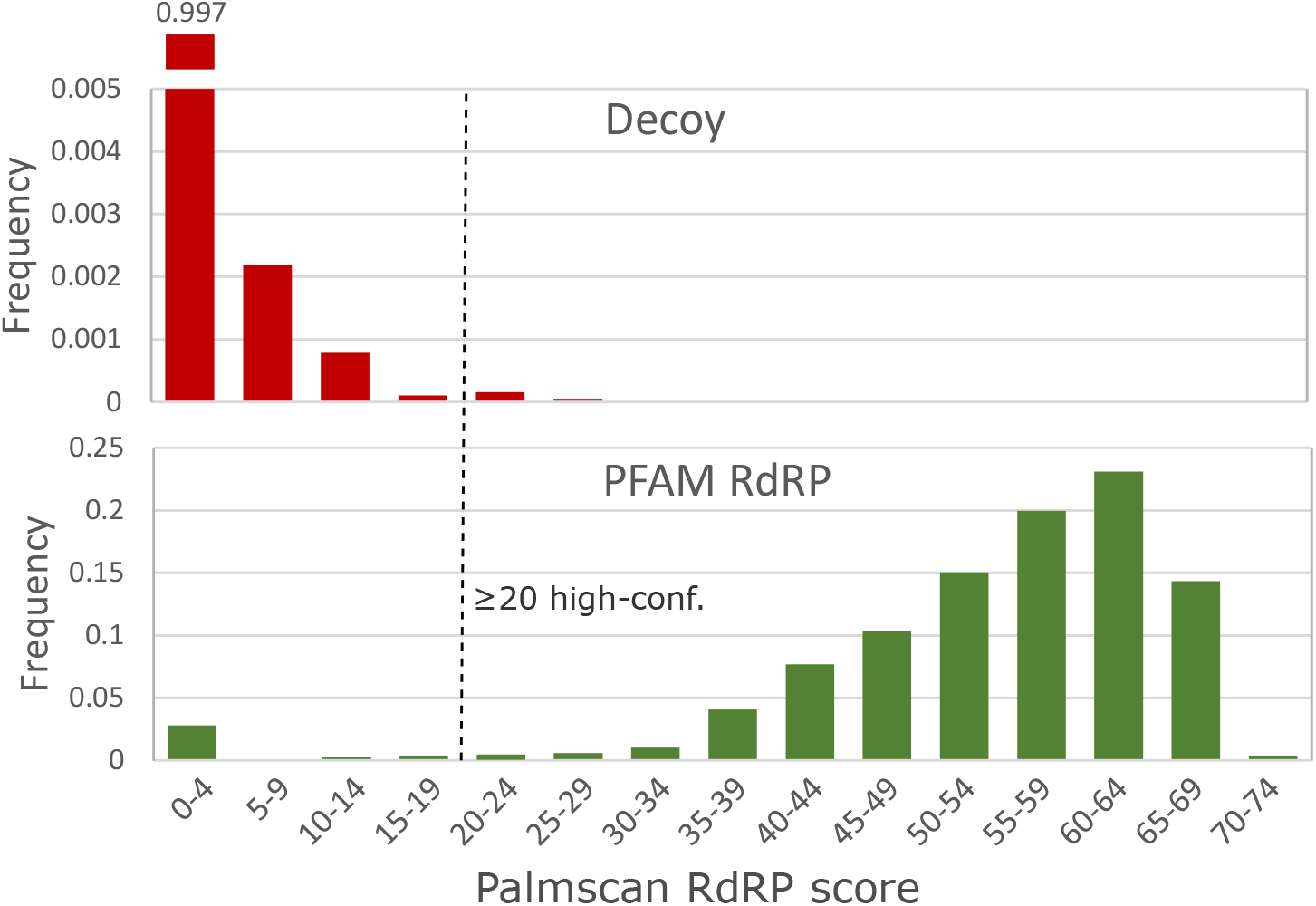
High-confidence palmprint score threshold. Distribution of RdRP palmprint scores on non-RdRP decoy set (top) and full PFAM RdRP alignments (bottom). The score threshold was set to 20 to discriminate RdRP polymerases with high confidence..

### OTU identity threshold tuning

We set the OTU clustering threshold to 90% after review of the results shown in Fig. 8, which shows that the number of OTUs is close to the number of species at this identity.

**Figure 8.**
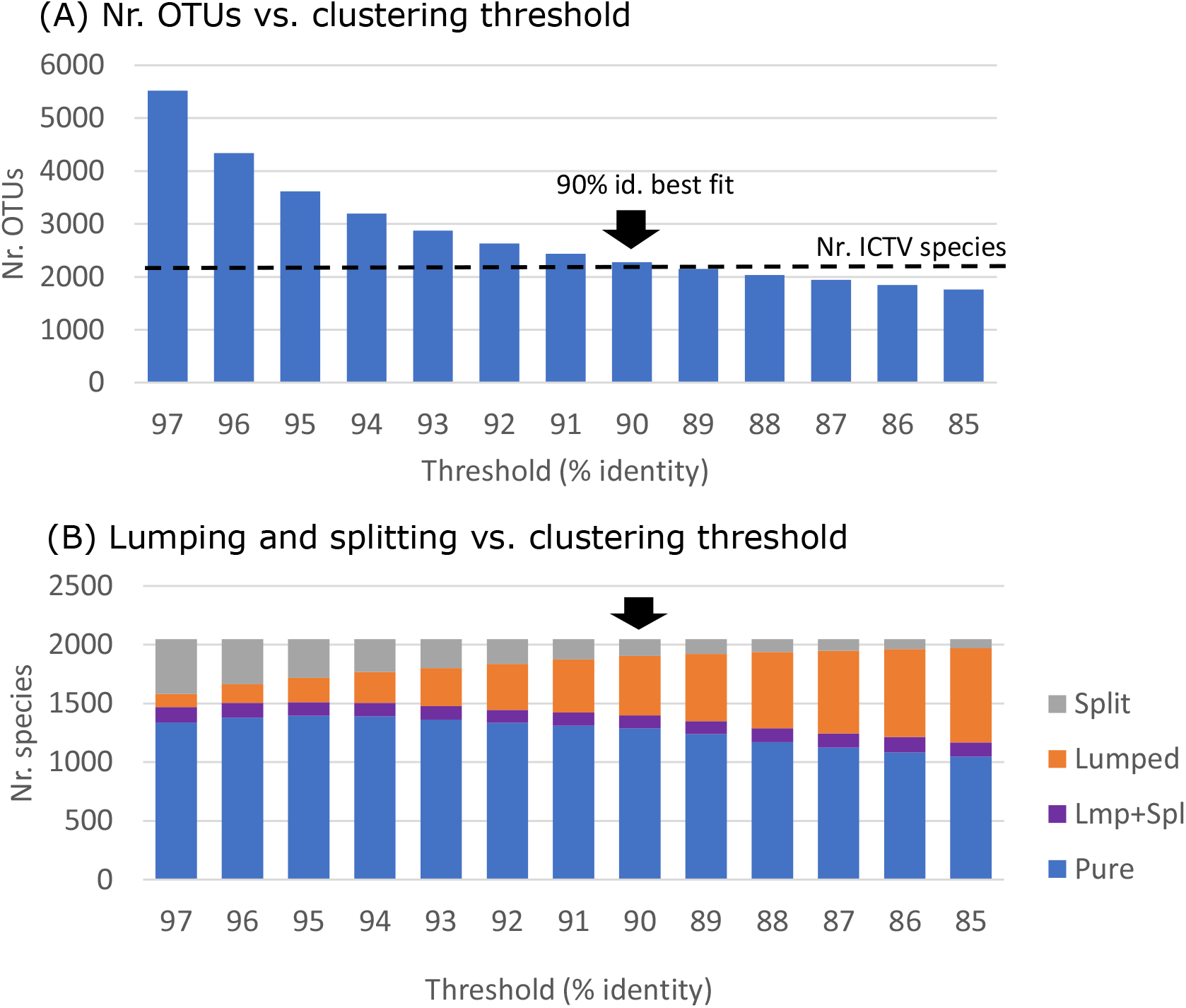
OTU identity threshold tuning. Top panel (A) shows the number of OTUs obtained by clustering RdRP palmprints of 2,048 recognised ICTV species at identity thresholds 97%, 96% … 85%. Lower panel (B) shows the number of species that are split over multiple OTUs, lumped together with one or more other species into a single OTU, both lumped and split (Lmp+Spl), or pure (not lumped or split). The best fit of number of clusters to number of species is obtained at 90% identity..

### Palmscan accuracy

Results on validation datasets are shown in Table 1. The sensitivity of `Palmscan`, measured as the fraction of positive sequences with a hit, is 87%. Combined, results on the the positive and negative datasets imply that on this data, the positive predictive value of `Palmscan` is 0.9989 and the false discovery rate is 0.0010. By contrast, `diamond` reports hits with E-value ≤ 10^−5^ in 88072/355449 = 27% of the non-RdRP sequences.

**Table 1.** Palmscan validation. *N* is the number of sequences in a dataset, *Pos/Neg* indicates whether the dataset is RdRP-positive (most or all sequences should contain RdRPs), or RdRP-negative (most sequences should not contain RdRP). *N* is the number of sequences, *N ps* is the number of `Palmscan` reported, and *N dmd* is the number of diamond hits with E-value ≤ 10^−5^. Notice the large numbers of diamond hits to non-RdRP sequences; in particular RTs. On the positive sets, `Palmscan` reports hits on a total of 3116*/*3572 = 87% of sequences.

| Set | Pos/Neg | N | N_ps | N_dmd |
| --- | --- | --- | --- | --- |
| PF00680_RdRP_1 | Pos | 795 | 760 | 795 |
| PF00978_RdRP_2 | Pos | 397 | 379 | 396 |
| PF00998_RdRP_3 | Pos | 205 | 194 | 204 |
| PF02123_RdRP_4 | Pos | 216 | 191 | 214 |
| Genomes | Pos | 1959 | 1592 | 1609 |
| PF00078_RT1 | Neg | 46876 | 0 | 13102 |
| PF07727_RT2 | Neg | 12037 | 0 | 1 |
| Decoy | Neg | 296536 | 4 | 74969 |

### Expect value of Palmscan false positives

`Palmscan` reported 64 palmprints on a random nucleotide sequence of length 10^11^ letters (100G) and 8,080 on a random amino acid sequence of the same length. The much higher false-positive rate on the amino acid sequence is primarily due to the absence of stop codons which often disrupt candidate palmprint sequences in the nucleotide case.

### Scope of PALMdb

PALMdb release `2021-03-02` contains 45,824 unique RdRP and RdRP-like palmprint sequences assigned to 15,016 OTUs.

## Discussion

### Implementing an RdRP barcode

Viral polymerases present challenges as a candidate for defining barcodes, including low sequence identity between diverged species, unclear domain boundaries, and permutations in the primary sequence of the active site where the best-conserved motifs are found. It is essential to identify barcode boundaries that can be reliably detected by automated sequence analysis to enable alignment between directly comparable sub-sequences of the gene. Otherwise, non-homologous regions may be included, creating the spurious appearance of greater evolutionary divergence, and/or fragments may overlap partially or be entirely disjoint, precluding estimates of evolutionary divergence from sequence alone. As an extreme example, disjoint fragments could belong to the same genome or to different families; if the only available information is disjoint sequence fragments, then no inferences can be made about their relatedness. Motifs A, B and C are present in most characterised viral polymerases, and defining a barcode as the minimal region including all amino acids in these motifs addresses the requirement for well-defined boundaries. However, automated identification of these boundaries is challenging due to low sequence identity and permutations. The `Palmscan` algorithm achieves high accuracy on this task, with good sensitivity of 87% on RdRP-containing sequences. The `Palmscan` false discovery rate was 0.0010 on a test set strongly focused on non-RdRP homologs of RdRP and other potentially RdRP-like sequences expected in metagenomic assemblies; while by contrast `diamond` reported hits to 27% of these decoy sequences with E-value *<* 10^*−*5^. Thus, `Palmscan` simultaneously performs well on two essential tasks in a high-throughput metagenomic analysis: (1) delineating boundaries for an informative barcode, and (2) discriminating RdRPs, which are very likely to be viral, from RTs which may be degraded host insertions.

### Barcode OTUs complement traditional taxonomic classification

The phenotype and pathogenecity of a virus is determined by its entire genome and interactions with its host cells. Functional changes in non-marker genes often accumulate more rapidly in non-polymerase genes. If a barcode sequence is identical to a well-characterised strain, then it is reasonable to infer that the phenotype is identical or similar; otherwise inferences become increasingly speculative as sequence identity diverges. Classification by barcode sequences is therefore unavoidably limited compared to careful examination of complete virus genomes and virion phenotype *in vitro* and *in vivo*. However, barcode classification is well suited to high-throughput analysis of large sequence datasets where metadata is limited and expert analysis is impractical. Thus, traditional taxonomy and barcode OTU classification approaches are complementary and mutually supporting.

### Defining species by an identity threshold

Within *Coronaviridae*, ICTV defines species as follows: “Viruses that share more than 90% aa sequence identity in the conserved replicase domains are considered to belong to the same species. This 90% identity threshold serves as the sole species demarcation criterion” (ICTV 2019b). This rule implicitly requires that identities of all pairs of coronaviruses must be considered, and aspires to a clustering where (a) all pairs within the same species have identity > 90% and (b) all pairs in different species have identity < 90%. However, criteria (a) and (b) cannot simultaneously be satisfied in practice, as shown by the examples in Fig. 9. Also, if this rule is strictly interpreted as written, then species must sometimes be merged as new viruses are discovered. Suppose that two viruses *X* and *Y* have identity < 90% and are assigned to different species. If a third virus *Z* is discovered which has identity *>* 90% to both *X* and *Y*, then according to the rule *Z* and *X* must belong to the same species and so must *Z* and *Y*. The only solution under a strict interpretation of the rule is to assign *X, Y* and *Z* to the same species, i.e. to merge the previously separate species of *X* and *Y*. This scenario is common in practice, as illustrated by Fig. 9. The ICTV rule corresponds to so-called single-linkage clustering where a node is joined to a cluster by any edge with distance less the threshold. An alternative rule is maximum linkage, where the distance to all other cluster members must be below the threshold. Maximum linkage clusters require existing species to be split into two or more species if a new virus is discovered which is > 90% to one existing member of the species but < 90% to another. Thus, both minimum and maximum linkage criteria are unstable by our definition, i.e. cause existing species assignments to change as new viruses are discovered. In our system, centroid clustering achieves stability by retaining existing centroids and adding new centroids for new viruses that fall outside of an existing cluster.

**Figure 9.**
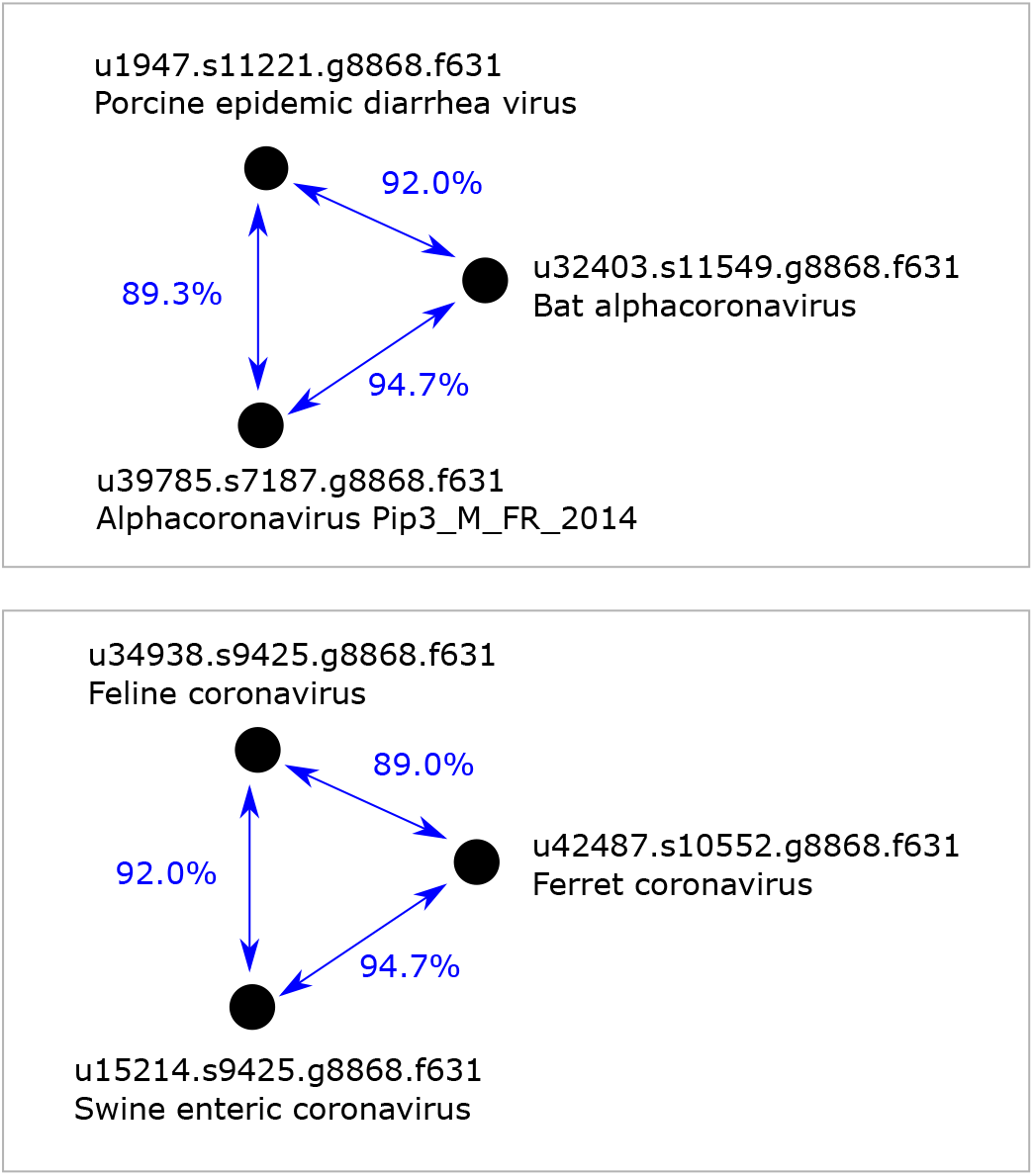
Examples of palmprint triples. In these examples, one out of three pairs has identity < 90% and two out of three pairs has identity > 90%. These examples show that it is not possible to define Coronavirus species such that all pairs in the same species have identity > 90% while all pairs in different species have identity < 90%.

### Centroid definition of species

The difficulty with the strict interpretation of the ICTV species definition can be resolved by interpreting existing species as centroids, as follows. If a newly-characterised virus *V* has ≥ 90% identity with a recognised virus, then it belongs to the same species, otherwise *V* is designated as the representative (centroid) of a new species.

### Sequence identity as a proxy for evolutionary divergence

The number of differences in a global alignment of two homologous sequences is an approximate indication of the total number of mutations since the most recent ancestral sequence. The number of inferred substitution mutations is underestimated if there are multiple substitutions at a single site, and the number of indel mutations is overestimated if a single event inserts or deletes multiple letters. Rearrangement events (inversions, tandem duplications, and translocations) disrupt global sequence homology, causing global alignment identity to overestimate divergence by forcing non-homologous segments to align. Permutation events in RdRP are translocations which disrupt global homology. `UCLUST` constructs global alignments, and will therefore underestimate sequence identity between palmprint sequences with different motif orders. An alternative approach is to design a method for arranging permuted sequences into canonical order, as implemented by (Wolf, Silas, et al. 2020). In our view, the homology of the variable regions V1 and V2 cannot always be reliably identified between permuted and canonically ordered domains. Also, an automated procedure for re-arranging permuted sequences would surely be subject to improvement, but any such improvements could cause instability by changing the OTU sequence of a known virus and hence its cluster assignment. We therefore chose to preserve the primary sequence of permuted palmprint sequences.

### Palmprints annotated as non-viral

Of the `Palmscan` hits to the source databases for OTU construction, 2, 749*/*36, 947 (7%) were annotated as non-viral. The majority of these are likely to be explained as endogenous viral elements (EVEs) or viral transcripts captured inadvertently in host-targeted sequencing projects and misassigned to the host. Curating these would require careful examination on a case-by-case basis, a considerable effort beyond the scope of the present work. We retain palmprints annotated as non-viral in our OTUs to aid in discriminating palmprints from viral genomes from palmprint-like sequences in host genomes, and because they offer opportunities for discovery of extant viruses for which similar sequences are otherwise unknown.

## Conclusions

We have shown that a segment of the viral polymerase palm domain, the palmprint, is well-suited for use as a taxonomic barcode. Our new algorithm for identifying this barcode, `Palmscan`, has good sensitivity to palmprints and is highly effective at discriminating RNA polymerases from reverse transcriptases. We provide PALMdb, an annotated reference database of RdRP palmprint sequences, to support sequence-based classification of riboviruses. We plan to extend PALMdb to include viral reverse transcriptases in the near future.

